# Secondary structural characterization of the nucleic acids from circular dichroism spectra using extreme gradient boosting decision-tree algorithm

**DOI:** 10.1101/2020.03.16.993352

**Authors:** Chakkarai Sathyaseelan, V Vinothini, Thenmalarchelvi Rathinavelan

## Abstract

Nucleic acids exhibit a repertoire of conformational preference depending on the sequence and environment. Circular dichroism (CD) is an important and valuable tool for monitoring such secondary structural conformations of nucleic acids. Nonetheless, the CD spectral diversity associated with these structures poses a challenge in obtaining the quantitative information about the secondary structural content of a given CD spectrum. To this end, the competence of extreme gradient boosting decision-tree algorithm has been exploited here to predict the diverse secondary structures of nucleic acids. A curated library of 610 CD spectra corresponding to 16 different secondary structures of nucleic acids has been developed and used as a training dataset. For a test dataset of 242 CD spectra, the algorithm exhibited the prediction accuracy of 99%. For the sake of accessibility, the entire process is automated and implemented as a webserver, called CD-NuSS (CD to nucleic acids secondary structure) and is freely accessible at https://www.iith.ac.in/cdnuss/. The XGBoost algorithm presented here may also be extended to identify the hybrid nucleic acid topologies in future.

## Introduction

Nucleic acids can take up a repertoire of secondary structural conformations such as right-handed double helix (B-form and A-type), left-handed double helix (Z-form), triplexes, quadruplexes, intercalated cytosine tetraplexes (i-motif) *etc*. These structures can either be inter or intramolecular in nature (1). They not only play a significant role in the regulation of various biological functions (2-4), but also, are responsible for several diseases (5,6). Like proteins (7), nucleic acids also exhibit optical activity (8) and each secondary structural conformations have distinct absorption for circularly polarized light (9,10). Circular dichroism (CD) spectroscopy is a simple, non-destructive optical technique that is most sensitive to the structural polymorphism of nucleic acids (1). Although CD cannot provide atomic-level structural insights about the nucleic acids like X-ray crystallography and NMR-spectroscopy, it has its own advantage in terms of quickness in providing the secondary structural information from a low sample concentration (7,11,12).

An accurate prediction of secondary structure and fold recognition of proteins from CD spectra is possible through various web servers (10,13-18). Similar to proteins, variations in the secondary structural architecture of nucleic acids affect their optical activity, thereby exhibiting a diverse CD spectra (1,3). Indeed, singular value decomposition (SVD) method has successfully been used in distinguishing the parallel, antiparallel and hybrid conformations of G-quadruplex (19). In line with this, machine learning algorithms have been employed here to identify 16 different secondary structures of nucleic acids (intra- and inter-molecular DNA and RNA), namely, A-form, antiparallel G quadruplex, antiparallel triplex, B-form, DNA RNA duplex, DNA stemloop, G-triplex, hybrid G quadruplex, iG pentaplex, iG quadruplex, i-motif, parallel G quadruplex, parallel duplex, parallel triplex, RNA stemloop and Z-form from the CD spectra.

The primary objective of any machine learning algorithm is to spot the hidden patterns or the trends in the data under examination, which is usually achieved by linear regression functions. These functions can be incorporated into feed forward neural network to segregate the data. One such single hidden layered feed-forward neural network that uses back propagation algorithm has previously been shown to predict five different secondary structures (helix, parallel and antiparallel β sheet, β turn, and random coil) of protein (20). Further, an optimized *Kohonen’s* self-organizing neural network map (21,22) (inspired from the sensorial nervous system of an animal) algorithm has been proved to accomplish protein topological (proteinotopic) mapping of the CD data to classify the proteins in a bi-dimensional map, and has been implemented as K2D web server (15,16,23). In fact, a similar attempt has also been made in the case of nucleic acids (21,22). As the nucleic acids can take up a variety of hybrid topologies such as hairpin-quadruplex (24-26), i-motif-quadruplex (27), triplex-duplex (24) *etc* to perform biological functions, it is essential to test the suitability of machine learning algorithms in predicting nucleic acids secondary structures. This may eventually be used in predicting the complex nucleic acids topologies when more training dataset corresponding to complex nucleic acids structures become available. Thus, the suitability of using three different supervised machine learning techniques *XGBoost, nnet* and *Kohonen* algorithms in characterizing the secondary structures of nucleic acids from the CD spectra has been investigated. The representative CD spectral features for different forms of nucleic acids are required to automate the aforementioned process. Thus, 610 CD spectral datasets heaped from the literature have been used as the training dataset for the above mentioned algorithms. The successfulness of the approach has been demonstrated by testing the trained model using 242 published CD spectral data. The results show that *XGBoost* method is superior in predicting the secondary structures of nucleic acids compared to the other two methods. The entire process has been automated and migrated into CD-NuSS (<u>c</u>ircular <u>d</u>ichroism to <u>nu</u>cleic acids <u>s</u>econdary <u>s</u>tructure) (https://www.iith.ac.in/cdnuss/) webserver, a tool that characterizes the secondary structures of nucleic acids using *XGBoost, Kohonen* and *nnet* machine learning algorithms.

## Materials and methods

The e<u>x</u>treme <u>g</u>radient <u>b</u>oosting (*XGBoost*) decision tree, <u>n</u>eural <u>n</u>etwork (*nnet*) and *Kohonen* super self-organized matrix (SOM) neural network algorithms have been considered in CD-NuSS webserver to predict 16 different forms of nucleic acids (**Figure 1a, Supplementary Figure S1**).

**Figure 1.**
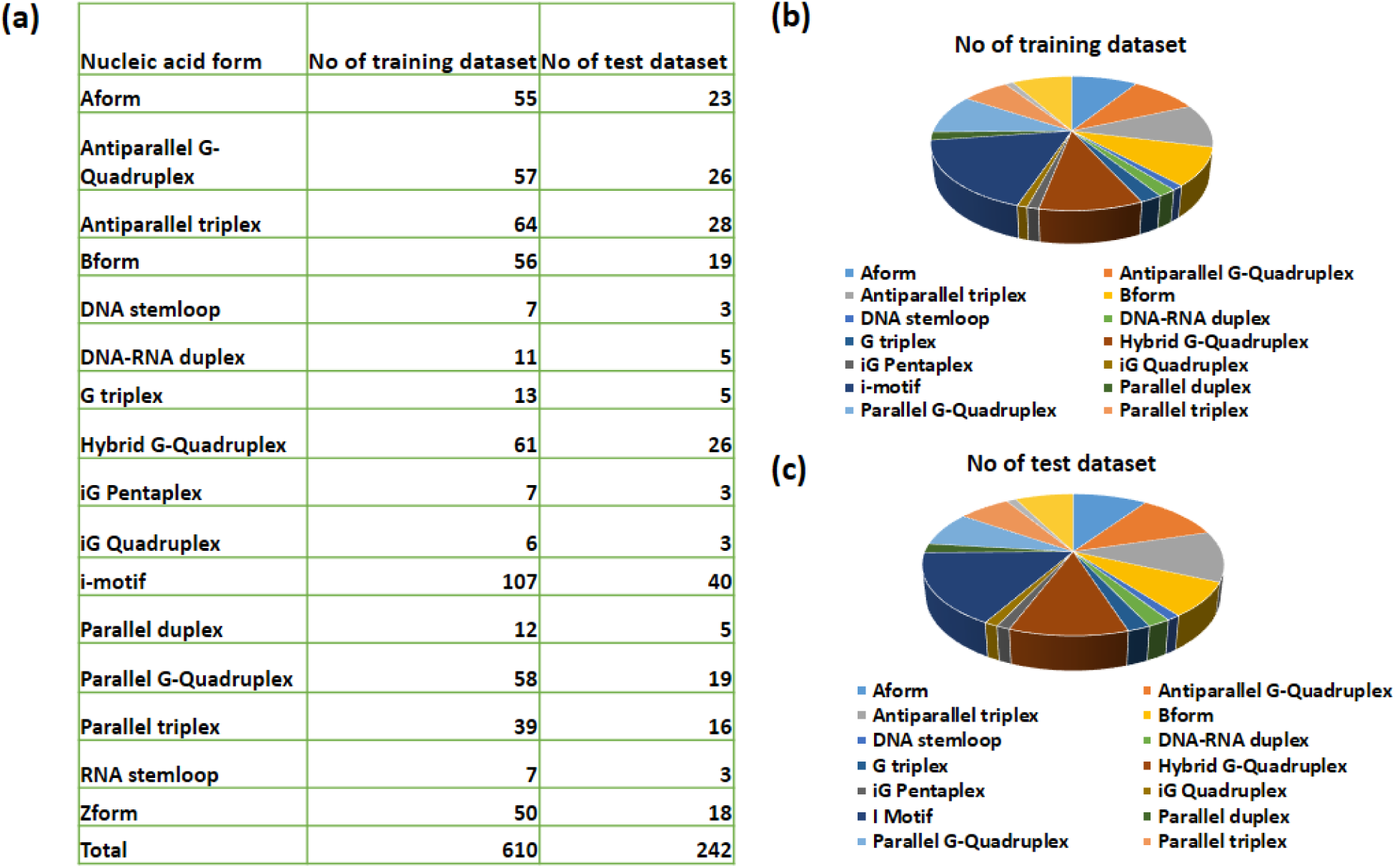
Statistics of the benchmark datasets used in the current investigation. (a) The number of training and test datasets used under 16 nucleic acids secondary structures and the corresponding pie chart illustrations of the (b) training and (c) test datasets.

### Benchmark dataset

A total of 852 CD spectral datasets have been collected through the literature survey **(Supplementary Table S1)**, among which, 610 and 242 datasets have been considered for training the model and testing the trained model respectively. The CD spectrum is ideally a plot between either ellipticity or molar ellipticity (Y-axis) against the wavelength (X-axis). However, instead of training the algorithm using the spectral image, numerical X- and Y-axes values are used here. Thus, the ellipticity or molar ellipticity in the CD spectra can be represented as an 86-dimensional vector corresponding to the wavelengths (1nm interval) in the range between 230 and 315nm.

The numerical data corresponding to the published CD spectral images were extracted using WebPlotDigitizer (28). Subsequently, a cubic spline algorithm has been employed to interpolate the numerical data points at a regular interval of 1nm from the CD spectral data points obtained from WebPlotDigitizer. Before feeding the data for training, the Y-axis data points corresponding to the 610 training datasets have been normalized in the range of -1.0 and +1.0nm using the formula given in the **equation (1)**.

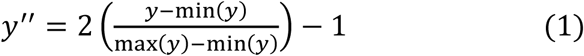

wherein, *y* and *y″* correspond to the ellipticity or molar ellipticity before and after normalization respectively and, min and max correspond to the minimum and maximum ellipticity or molar ellipticity respectively. This step has been performed to create a bi-dimensional map that retain the topology of the spectra corresponding to different forms of nucleic acids by ignoring the impact of the experimental conditions.

### Implementation of CD-NuSS

Few successful attempts have been made to predict the secondary structures of proteins using the supervised machine learning (15,16,21-23). Here, the existing *Kohonen (21,22), nnet (29)* and *XGBoost (30)* modules in R-package (*Kohonen* and *nnet*) or python (*XGBoost*) were used as an template and modified accordingly to train the model and test the sample dataset.

### Prediction of the nucleic acids secondary structures using eXtreme Gradient Boosting (XGBoost) algorithm

The *XGBoost* algorithm uses the gradient boosting machine (GBM) framework, wherein, the tree ensemble (a set of classification and regression trees (CART)) is boosted through a sequential learning. **Figure 2a** illustrates the flow of the sequential boosting process involved in the training of 16 different nucleic acids conformations. As the *XGBoost* works only with the numerical vectors, the A-form, antiparallel G quadruplex, antiparallel triplex, B-form, DNA RNA duplex, DNA stemloop, G-triplex, hybrid G quadruplex, iG pentaplex, iG quadruplex, i-motif, parallel G quadruplex, parallel duplex, parallel triplex, RNA stemloop and Z-form from the CD spectra are labeled as 0, 1, 2, 3, 4, 5, 6, 7, 8, 9, 10, 11, 12, 13, 14 and 15 respectively. The learning objective is set as *multi:soft prob* to perform multi-class classification (16 nucleic acids classifiction), wherein, the output vector is reshaped into 86*16 (ndata*nclass) matrix. The aforementioned matrix contains the predicted probability of each data point (86 data points corresponding to 86 wavelength) belonging to each class (16 secondary structures). During the training, the rate of learning has been set as 0.1 step size shrinkage (as it slowdowns the learning) to prevent overfitting and the boosting has been done on an ensemble of 500 trees to model the training dataset.

**Figure 2.**
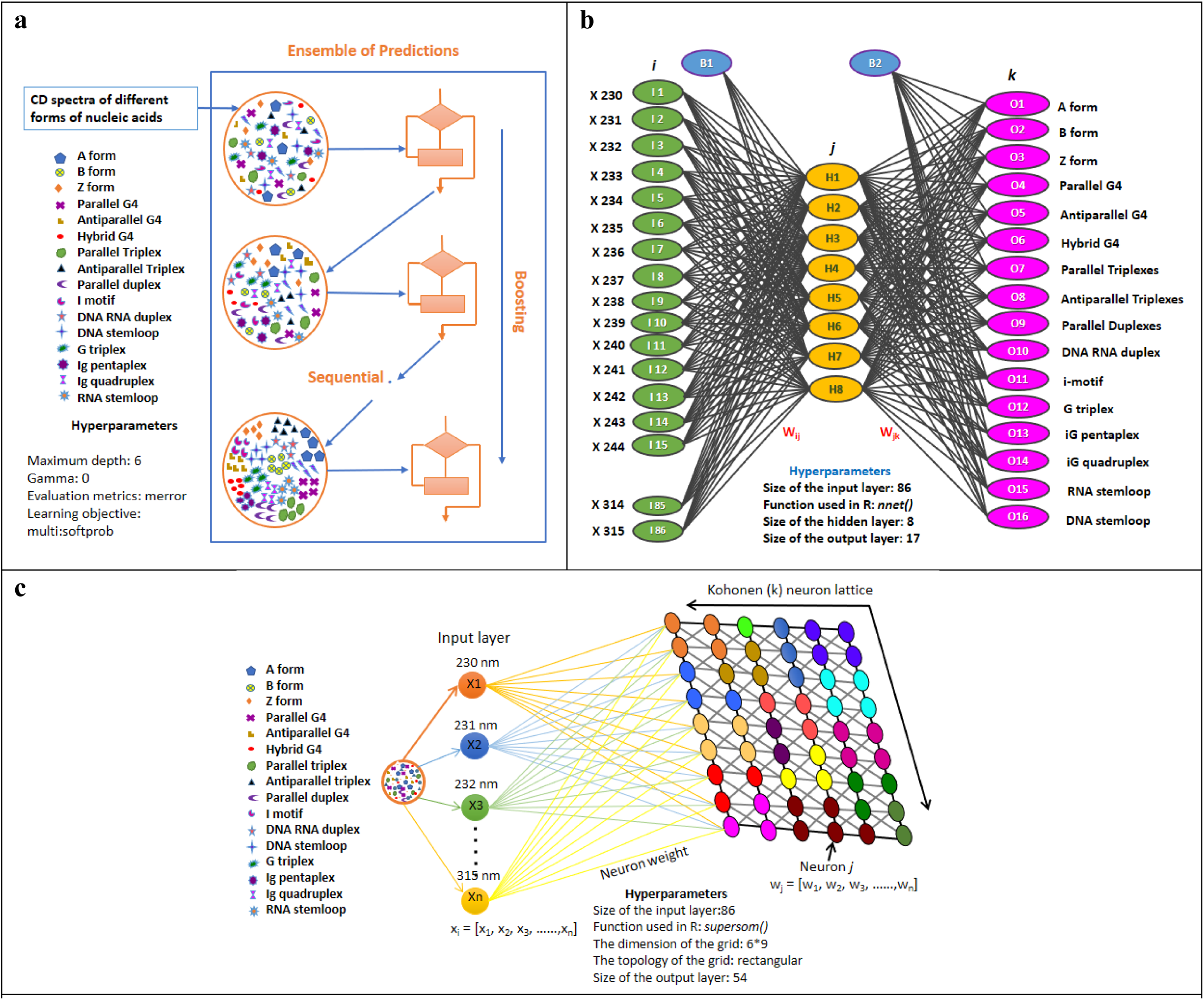
**The schematic illustration of: (a)** eXtreme gradient boosting (*XGBoost*) algorithm used in the prediction process depicts the logic in which the subsequent predictors learn sequentially from the mistake of the previous prediction. A list of hyperparameter values used for *XGBoost* method are maximum depth:6, gamma:0, evaluation metrics: merror and learning objective: multi:softprob. (**b)** feed-forward neural network algorithm used in training the 610 CD samples corresponding to 16 different nucleic acids secondary structures. Green, yellow and purple circles represent the input (I1 to I86), hidden (H1 to H8) and output (O1 to O16) layer respectively. B1 and B2 are the bias nodes. X230 to X315 indicate the ellipticity values corresponding to different nucleic acids in the wavelength range between 230 and 315nm. W_ij_ and W_jk_ being the weight of neurons from the input layer (i) to the hidden layer (j) and from the hidden layer (j) to the output layer (k) respectively. A list of hyperparameter values used for *nnet* method are *nnet()* function, 8 as hidden layer size and 17 as output layer size. **(c)** *Kohonen* self-organizing map with 86 input vectors indicated as x1, x2, x3,…x86 correspond to the wavelengths in the range of 230 to 315nm in 1nm interval in the CD spectra. *Kohonen* 2D lattice has 54 neurons and W_j_ is the weight matrix of the neurons w1,w2,w3,…,w54. A list of hyperparameter values used for the *Kohonen* method are supersom() function, 6*9 grid dimension, rectangular grid topology *etc.* See text for details.

### Prediction of the nucleic acids secondary structures using the neural network algorithms nnet algorithm

In the feed-forward neural network (*nnet*) algorithm, there are 86 input vectors (representing the wavelengths in the range of 230 to 315nm in 1nm interval in the CD spectra) in the input layer. Input layer ‘fans out’ and distributes the input vectors to the hidden units in such a way that each input vector is connected to each of the 8 units in the hidden layer (**Figure 2b**). The hidden neurons of 8 are chosen here as they are found to be optimal for the training dataset used here to give a better performance of the algorithm. W_ij_ is the weight matrix that connects the input vector and the neurons in the hidden layer and is assigned random value initially and optimized in the subsequent rounds. The neurons in the hidden layer takes a fixed activation function ϕ_j_ summing up the values in the hidden units and a bias constant (B1). A default value of 1 is used for B1 here and is optimized based on the values of input layer. Similarly, the output activation function ϕ_o_ is the summation of the hidden units and a bias constant (B2). A default value of 1 is used for B2 here and is optimized based on the values of hidden layer. The 8 units in the hidden layer “fans out” to 16 different output units in the output layer, representing 16 different nucleic acids secondary structures. For a unit *j* in a hidden or output layer, the net input *I*_*j*_ to the unit *j* is given in equation (2).

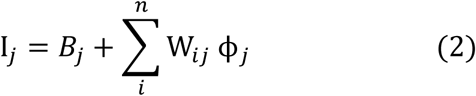

wherein, B_j_ is the bias constant, W_ij_ is weight between input i and output j, and ϕ_*j*_ is the activation function

Given the net input *I*_*j*_ to unit *j*, then the output of the unit *j* (O_*j*_) is computed by equation (3). This (O_*j*_) function is also called as squashing function, because it maps a large input domain onto the smaller range of 0 to 1

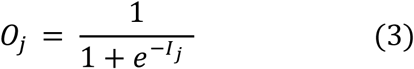

### Kohonen algorithm

The input layer of the *Kohonen* self-organizing map (SOM) is a fully connected neural network with ‘n’ units (*viz.*, the length of the training vector), wherein, ‘n’ is 86 here that corresponds to the wavelengths in the CD spectra (230 to 315nm at 1nm interval). Each training input unit has the ellipticity or molar ellipticity values corresponding to 16 different secondary structures of nucleic acids from multiple datasets. The input layer is fully connected to the *Kohonen* 2D lattice that is of 6×9 dimension (totally 54 neurons) through a randomly assigned weight matrix W_j_ (wherein, ‘j’ stands for the number of neurons in the *Kohonen* lattice) (**Figure 2c**). It is noteworthy that 54 *Kohonen* neurons are chosen as it gives better secondary structure prediction accuracy (for the test dataset) using the training dataset considered here. The learning rate (α) is set to 0.05 (chosen based on the prediction accuracy), and the training continues until the rate converges. The training of the model has been done in an iterative fashion for 86 cycles (as there are 86 wavelengths between 230-315nm at 1nm interval), during which, the weight matrix is updated by giving more weightage to the neuron (winning neuron) that has a shorted Euclidian distance (between input vector and weight vector.

### Automation

The algorithms described above for predicting the secondary structures of 16 different nucleic acids conformations and estimating the K_d_ have been transformed into an automated webserver that runs on Ubuntu Linux (18.04 LTS). The interactive web server has been developed with Apache (https://httpd.apache.org) and D3.js (https://d3js.org/). The client-side user interface has been implemented using HTML and PHP. The webserver, namely, CDNuSS (<u>c</u>ircular <u>d</u>ichroism to <u>n</u>ucleic <u>a</u>cids <u>s</u>econdary <u>s</u>tructure) is freely available at (https://www.iith.ac.in/cdnuss/).

## Results and discussion

### Topology of the CD spectra corresponding to different nucleic acids secondary structures

A close inspection of the training set corresponding to 16 different nucleic acids conformations indicate that these conformations have signature peaks, thus, are amicable for machine learning algorithms. For instance, the negative and positive peaks around 245-250nm and 275-280nm respectively represent the B-form. The i-motif has a negative peak around 265nm and a positive peak around 285-290nm, the antiparallel triplex has a negative peak around 245nm and a positive peak around 280nm and the parallel triplex has a negative peak around 255nm and a positive peak around 290nm **(Supplementary Table S2)**, *etc*.

### Functionality of the CD-NuSS web server

Upon establishing the methodology (discussed under Methods), the entire process is automated and migrated into a web interface, namely CD-NuSS. The **Figure 3** depicts the web interface, wherein, the user can upload the CD spectral information (in text format) and choose among the *XGBoost, nnet* and *Kohonen* algorithms to characterize the nucleic acids secondary structure.

**Figure 3.**
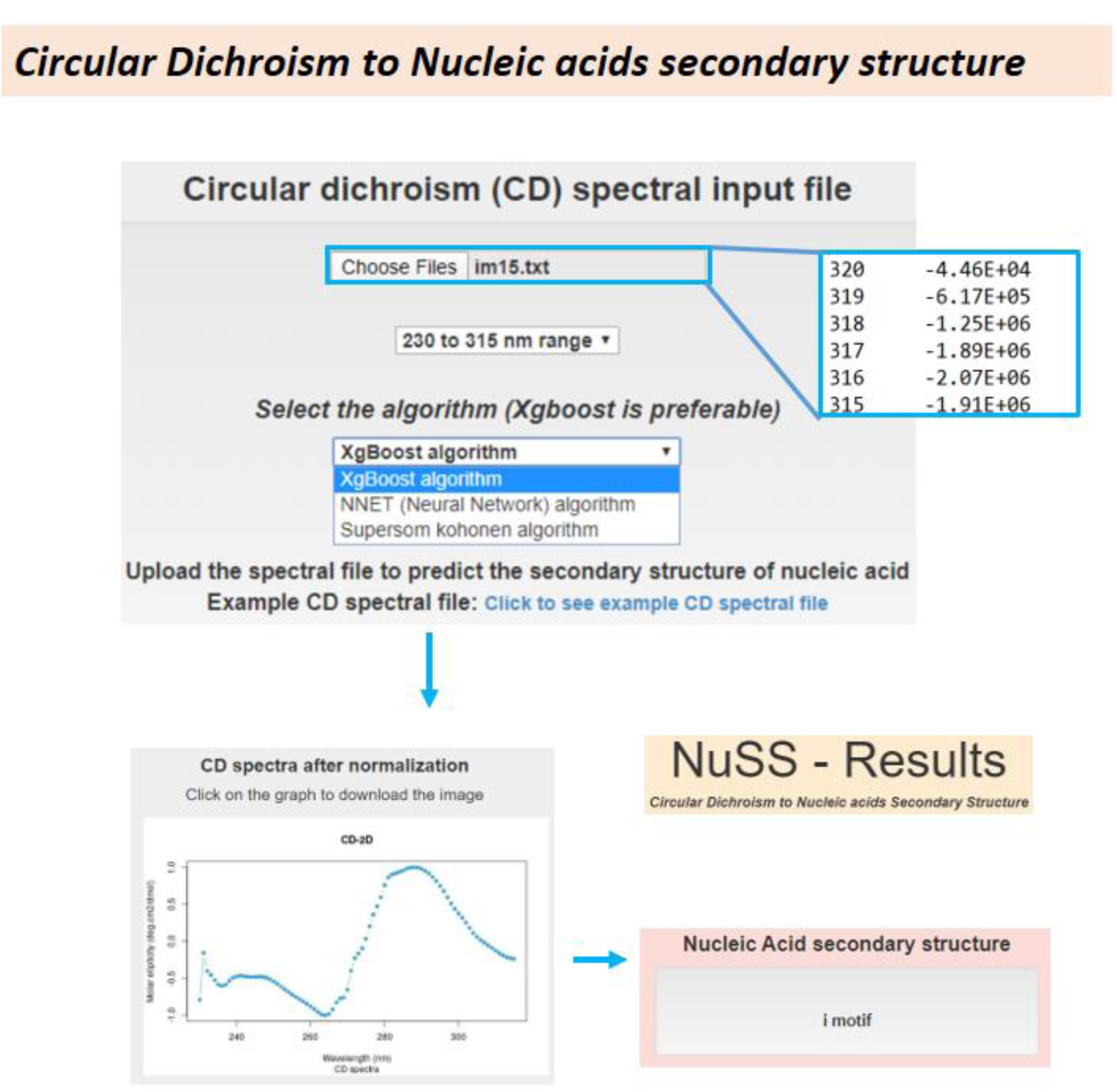
CD-NuSS web interface. The snapshots describing the nucleic acid secondary structure prediction. Note that the predicted secondary structure shown at the bottom. See text for details.

### Automated characterization of the nuclei acids secondary structures

Following the training of the 610 samples using *XGBoost, Kohonen* and *nnet* and algorithms, the efficacy of these algorithms in predicting the secondary structures of the nucleic acids has been tested through the server. By considering 242 test samples (**Figure 1a, 1c**), the efficacy of the *XGBoost, nnet* and super-SOM *Kohonen* algorithms in predicting the secondary structures of the nucleic acids has been tested in CD-NuSS web server. The server accepts the text format input file that has the ellipticity or molar ellipticity values against the wavelength. Before the prediction, the test samples are normalized using the **equation 1** to a common scale in the Y-axis such a way that the ellipticity falls in the range of -1.0 and +1.0 while retaining the topology of the spectrum **(Supplementary Figure S2).** This improves the prediction accuracy by circumventing the effect of sample conditions. It is noteworthy that the reference training dataset is also normalized prior to the training as discussed in the Methods section. As the published CD spectra are mainly in the wavelength region of 230 to 315nm, the training and prediction is based only on this wavelength region.

### XGBoost decision tree algorithm exhibits higher prediction accuracy compared with Kohonen and nnet

The **Supplementary Table S1** represents the 16*16 confusion matrix, which reflect the accuracy of the secondary structure prediction of nucleic acids using *XGBoost* algorithm. Out of the 242 test samples, 2 false positives have been observed. For instance, a B form conformation is falsely predicted as an antiparallel triplex and a parallel G quadruplex is falsely predicted as an i-motif conformation as their signature peaks falls broadly under the false positive categories. Overall, the *XGBoost* algorithm provides 99% prediction accuracy. As can be seen in **Supplementary Table S3**, the 16*16 confusion matrix corresponding to the *nnet* algorithm has 43 false positives followed by the *Kohonen* algorithm which has 86 false positives. These are comparatively higher than the *XGBoost* algorithm. Out of 242 test sets, *XGBoost, nnet* and *Kohonen* algorithms have accurately predicted 240, 199 and 156 secondary structures respectively **(Supplementary Table S4).** Thus, the order of prediction accuracy is: *XGBoost* > *nnet* > *Kohonen* with an overall accuracy of 99%, 82% and 64% respectively. **Figure 4** illustrate the success rate comparison of *XGBoost, nnet* and *Kohonen* algorithms in predicting the 16 different nucleic acids secondary structures.

**Figure 4.**
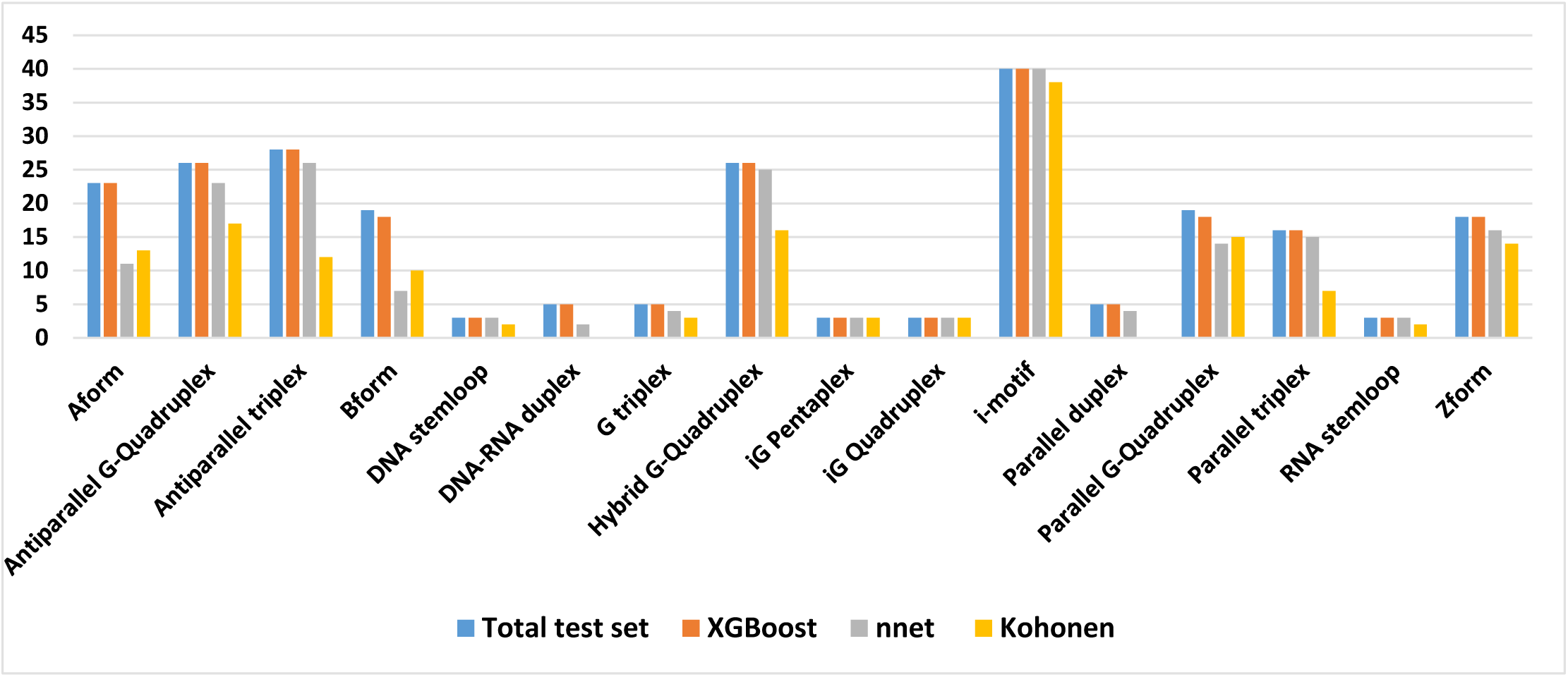
Histogram illustrating the secondary structural prediction accuracy of *Kohonen, nnet* and *XGBoost* algorithms for the test datasets (Figure 1c). Note that the prediction accuracy is higher for the *XGBoost* decision tree algorithm compared with the other two methods

## Conclusion

The efficacy of *XGBoost* decision tree algorithm, *nnet* and feed forward (super self-organizing map) *Kohonen* neural network algorithms in predicting the 16 different secondary structures of nucleic acids have been investigated in the current study. The reference dataset has been initially trained by considering 610 training samples for each algorithm and 242 sample datasets have been used for the testing. The results indicate that *XGBoost* method is superior among all the other methods with 99% prediction accuracy and the *Kohonen* algorithm has the lowest prediction accuracy of 64%.

## Acknowledgements

The authors thank IITH and CDAC for the computational resources. VV and CS thank Ministry of Human Resource Development, Government of India for the fellowships. The work was supported by BIRAC-SRISTI GYTI award (PMU_2017_010), BIRAC-SRISTI GYTI award (PMU_2019) and Indian Institute of Technology Hyderabad (IITH).

## Author contributions

TR designed and supervised the project. VV collected the sample sets and developed the methodology in R. CS improved the sample size, fine-tuned the prediction efficiency and implemented the web interface. VV, CS and TR wrote the manuscript.

## Conflict of interest

None

## Reference

1. Kypr, J., Kejnovska, I., Renciuk, D. and Vorlickova, M. (2009) Circular dichroism and conformational polymorphism of DNA. Nucleic acids research, 37, 1713–1725.

2. Koster, D.A., Crut, A., Shuman, S., Bjornsti, M.A. and Dekker, N.H. (2010) Cellular strategies for regulating DNA supercoiling: a single-molecule perspective. Cell, 142, 519–530.

3. J. Kypr, I. Kejnovska and Vorlìcková, K.B.a.M. (2012) Circular Dichroism Spectroscopy of Nucleic Acids in Comprehensive Chiroptical Spectroscopy. John Wiley & Sons, Inc., Hoboken, NJ, USA.

4. G. D. Fasman. (2001) Circular Dichroism and the Conformational Analysis of Biomolecules. Springer-Verlag, New York, NJ, USA.

5. Herbert, A. (2019) Z-DNA and Z-RNA in human disease. Communications biology, 2, 7.

6. Zhou, B., Geng, Y., Liu, C., Miao, H., Ren, Y., Xu, N., Shi, X., You, Y., Lee, T. and Zhu, G. (2018) Characterizations of distinct parallel and antiparallel G-quadruplexes formed by two-repeat ALS and FTD related GGGGCC sequence. Scientific reports, 8, 2366.

7. Greenfield, N.J. (2006) Using circular dichroism spectra to estimate protein secondary structure. Nature protocols, 1, 2876–2890.

8. Tinoco, I., Jr., Bustamante, C. and Maestre, M.F. (1980) The optical activity of nucleic acids and their aggregates. Annual review of biophysics and bioengineering, 9, 107–141.

9. Drake, A.F. (1994) Circular dichroism. Methods in molecular biology, 22, 219–244.

10. Micsonai, A., Wien, F., Kernya, L., Lee, Y.H., Goto, Y., Refregiers, M. and Kardos, J. (2015) Accurate secondary structure prediction and fold recognition for circular dichroism spectroscopy. Proceedings of the National Academy of Sciences of the United States of America, 112, E3095–3103.

11. Miles, A.J. and Wallace, B.A. (2016) Circular dichroism spectroscopy of membrane proteins. Chemical Society reviews, 45, 4859–4872.

12. Wallace, B.A. (2019) The role of circular dichroism spectroscopy in the era of integrative structural biology. Current opinion in structural biology, 58, 191–196.

13. Micsonai, A., Wien, F., Bulyaki, E., Kun, J., Moussong, E., Lee, Y.H., Goto, Y., Refregiers, M. and Kardos, J. (2018) BeStSel: a web server for accurate protein secondary structure prediction and fold recognition from the circular dichroism spectra. Nucleic acids research, 46, W315–W322.

14. Xue, L.C., Rodrigues, J.P., Kastritis, P.L., Bonvin, A.M. and Vangone, A. (2016) PRODIGY: a web server for predicting the binding affinity of protein-protein complexes. Bioinformatics, 32, 3676–3678.

15. Perez-Iratxeta, C. and Andrade-Navarro, M.A. (2008) K2D2: estimation of protein secondary structure from circular dichroism spectra. BMC structural biology, 8, 25.

16. Andrade, M.A., Chacon, P., Merelo, J.J. and Moran, F. (1993) Evaluation of secondary structure of proteins from UV circular dichroism spectra using an unsupervised learning neural network. Protein engineering, 6, 383–390.

17. Whitmore, L. and Wallace, B.A. (2008) Protein secondary structure analyses from circular dichroism spectroscopy: methods and reference databases. Biopolymers, 89, 392–400.

18. Whitmore, L. and Wallace, B.A. (2004) DICHROWEB, an online server for protein secondary structure analyses from circular dichroism spectroscopic data. Nucleic acids research, 32, W668–673.

19. Del Villar-Guerra, R., Trent, J.O. and Chaires, J.B. (2018) G-Quadruplex Secondary Structure Obtained from Circular Dichroism Spectroscopy. Angewandte Chemie, 57, 7171–7175.

20. Kaur, H. and Raghava, G.P. (2003) Prediction of beta-turns in proteins from multiple alignment using neural network. Protein science : a publication of the Protein Society, 12, 627–634.

21. R, W. and J, K. (2018) Flexible Self-Organizing Maps in kohonen 3.0. Journal of Statistical Software, 87, 1–18.

22. R, W. and Lmc, B. (2007) Self- and Super-Organizing Maps in R: The kohonen Package. Journal of Statistical Software, 21, 1–19.

23. Bethapudi, S. and Desai, S. (2018) Separation of pulsar signals from noise using supervised machine learning algorithms. Astronomy and Computing, 23, 15–26.

24. Hou, X.M., Fu, Y.B., Wu, W.Q., Wang, L., Teng, F.Y., Xie, P., Wang, P.Y. and Xi, X.G. (2017) Involvement of G-triplex and G-hairpin in the multi-pathway folding of human telomeric G-quadruplex. Nucleic acids research, 45, 11401–11412.

25. Lim, K.W., Khong, Z.J. and Phan, A.T. (2014) Thermal stability of DNA quadruplex-duplex hybrids. Biochemistry, 53, 247–257.

26. Lim, K.W. and Phan, A.T. (2013) Structural basis of DNA quadruplex-duplex junction formation. Angewandte Chemie, 52, 8566–8569.

27. Chu, B., Zhang, D. and Paukstelis, P.J. (2019) A DNA G-quadruplex/i-motif hybrid. Nucleic acids research, 47, 11921–11930.

28. Rohatgi, A. (2019) WebPlotDigitizer - Version 4.2. KDD ‘16 Proceedings of the 22nd ACM SIGKDD International Conference on Knowledge Discovery and Data Mining.

29. Venables, W.N. and Ripley, B.D. (2002) Modern Applied Statistics with S. Springer, XII, 498.

30. Chen, T. and Guestrin, C. (2016) XGBoost: A Scalable Tree Boosting System. KDD ‘16 Proceedings of the 22nd ACM SIGKDD International Conference on Knowledge Discovery and Data Mining, 785–794

