## supplementary file for "Secondary structural characterization of the nucleic acids from circular dichroism spectra using extreme gradient boosting decision-tree algorithm"

**Title**

### Supplementary Figures

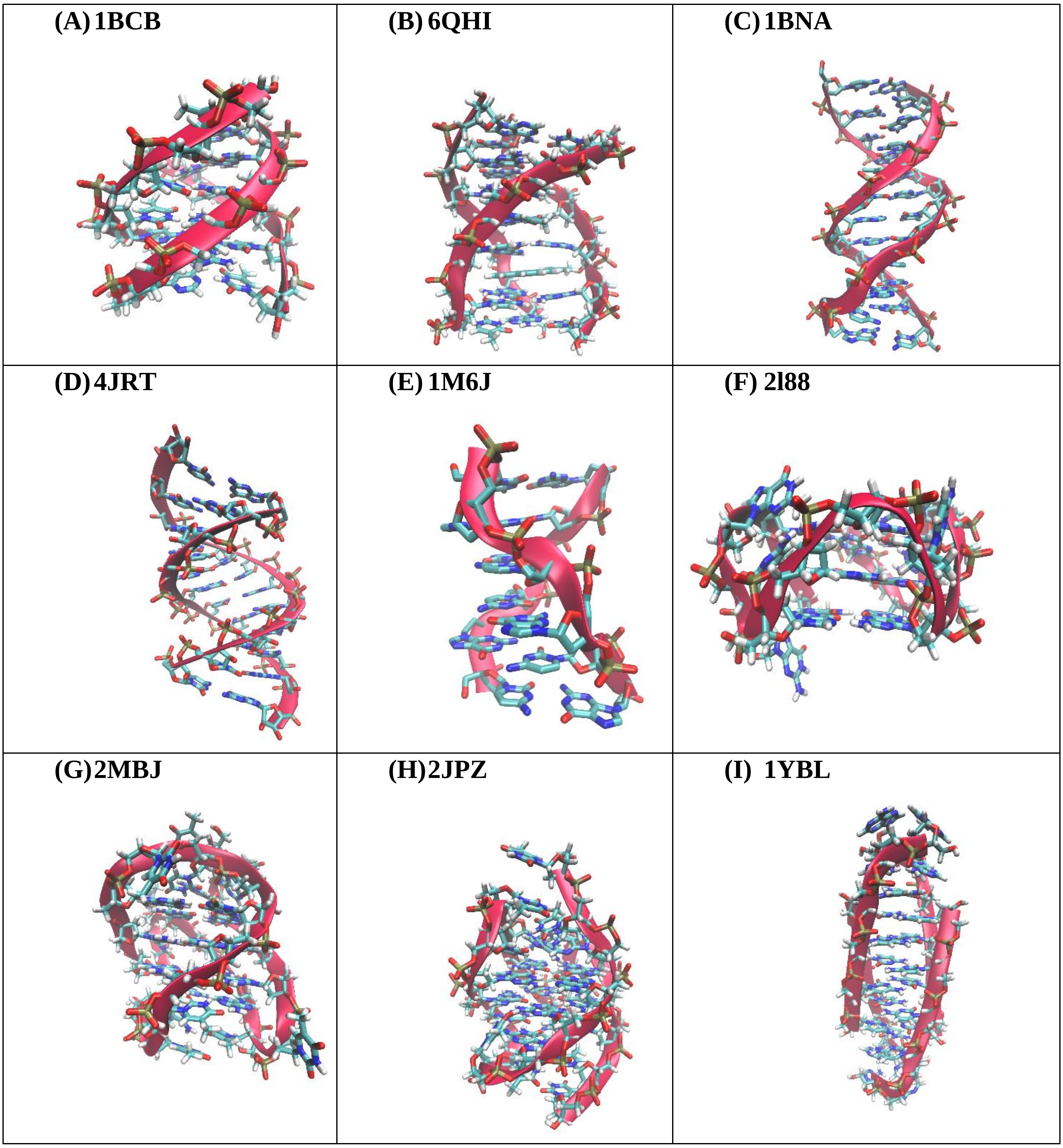

**(J) 2LUJ**

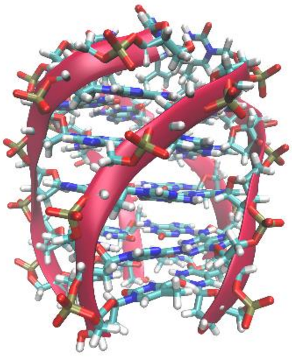

**(K) 2MKO**

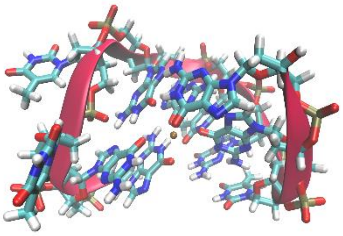

**(L) 1WKS**

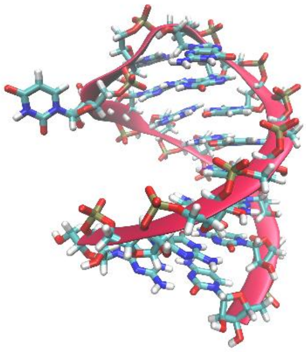

**(M) 1EFS**

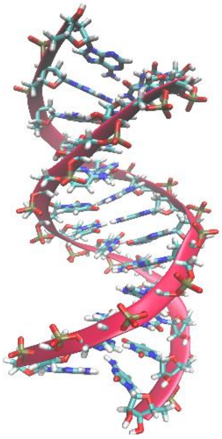

**(N) 1R2L**

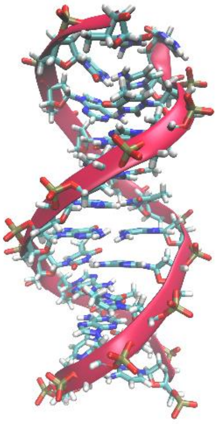

**(O) 1OLD**

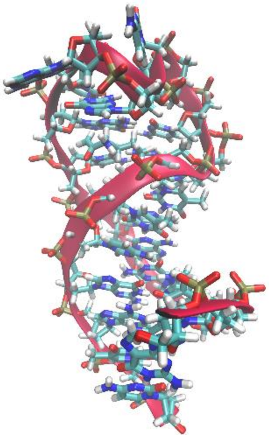

**Figure S1. Cartoon representation of different forms of nucleic acids secondary structures considered in the current investigation:** A) parallel triplex B) antiparallel triplex C) Bform DNA D) Aform RNA E) Zform F) parallel G quadruplex G) antiparallel G quadruplex H) hybrid G Quadruplex I) i-motif J) iG pentaplex K) G triplex L) RNA stemloop M) DNA RNA duplex N) parallel duplex O) DNA stemloop. VMD (http://www.ks.uiuc.edu/Research/vmd/) is used to generate the cartoon representation. Note that the atomic structure corresponding to iG quadruplex conformation is not yet available.

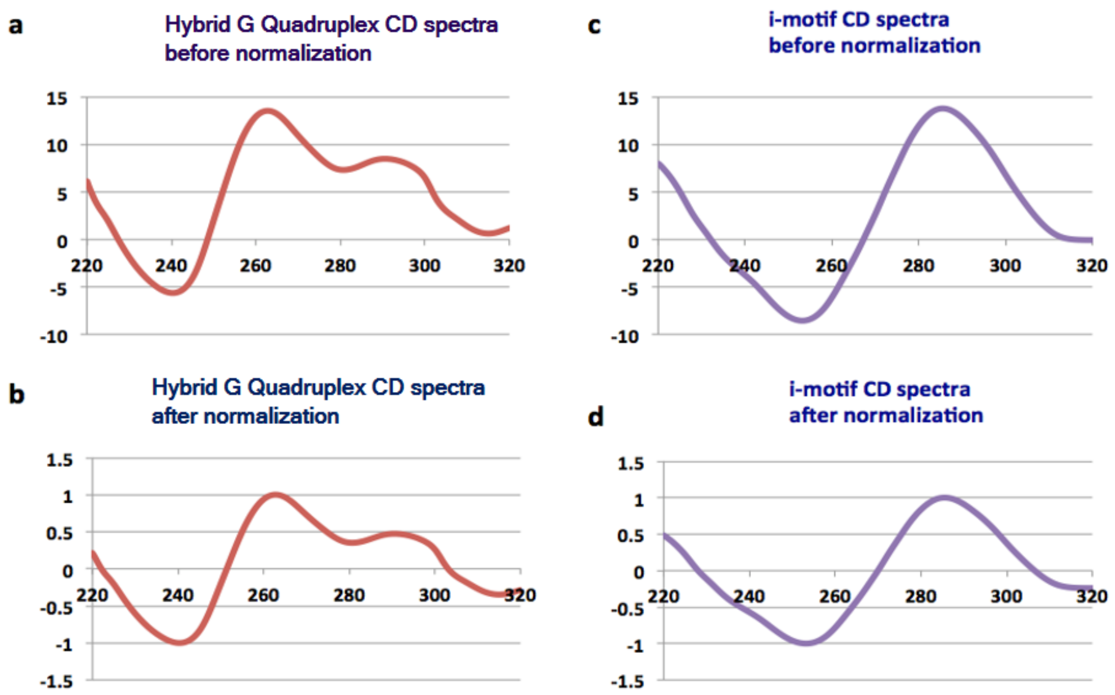


**Figure S2. Representative CD spectral graphs showing the preservation of the spectral topology after normalization.** The CD spectra corresponding to the Hybrid G-Quadruplex ((a) before and (b) after normalization) and i-motif ((c) before and (d) after normalization). Note that the signature peaks of the Hybrid G-quadruplex (a negative peak at 240nm, a positive peak at ~260nm and a slight positive peak ~290nm) and the i-motif (a negative peak at ~250nm and a positive peak at ~280-285nm) are retained.

**Supplementary Tables**

**Table S1. The references from which the CD spectra has been collected and used for the training and testing.**

| S.No | Nucleic acid form | Reference DOI number |
|---|---|---|
| 1 | Aform-DNA | 10.1093/nar/gkp026 |
| 2 | Aform-DNA | 10.1111/j.1747-0285.2009.00847.x |
| 3 | Aform-DNA | 10.1016/0076-6879(95)46006-3 |
| 4 | Aform-DNA | 10.1093/nar/gkl513 |
| 5 | Aform-DNA | Darwish, Maged A., "Stability of Nucleic Acid Secondary Structures and Their Contribution to Gene Expression" (2010). *Seton Hall University Dissertations and Theses (ETDs)*. 1435. |
| 6 | Aform-RNA | 10.1093/nar/gkp026 |
| 7 | Aform-RNA | 10.1002/chir.22064 |
| 8 | Aform-RNA | 10.1111/j.1747-0285.2009.00847.x |
| 9 | Aform-RNA | 10.1002/0471142700.nc0711s11 |
| 10 | Aform-DNA | 10.1002/chir.22064 |
| 11 | Aform-DNA | 10.1093/nar/gkl513 |
| 12 | Aform-DNA | 10.3390/molecules23102572 |
| 13 | Aform-DNA | 10.1002/bip.1979.360180418 |
| 14 | Aform-DNA | 10.1093/nar/13.13.4983 |
| 15 | Aform-RNA | 10.3390/molecules23102572 |
| 16 | Aform-RNA | 10.1016/j.bpj.2016.11.018 |
| 17 | Aform-RNA | 10.1038/s41598-019-40715-2 |
| 18 | Bform | 10.1073/pnas.94.16.8421 |
| 19 | Bform | 10.1016/j.biomaterials.2004.07.038 |
| 20 | Bform | 10.1093/nar/gkp026 |
| 21 | Bform | 10.1002/9781118120392.ch17 |
| 22 | Bform | 10.1016/s0304-4165(00)00063-5 |
| 23 | Bform | 10.1073/pnas.0911528107 |
| 24 | Bform | 10.1002/chir.22064 |

| | | |
|---|---|---|
| 25 | Bform | 10.1093/nar/29.13.2795 |
| 26 | Bform | 10.1111/j.1747-0285.2009.00847.x |
| 27 | Bform | 10.1038/365566a0 |
| 28 | Bform | 10.1073/pnas.78.6.3546 |
| 29 | Bform | 10.1371/journal.pone.0198418 |
| 30 | Bform | 10.1021/jp3085556 |
| 31 | Bform | 10.1093/nar/gkl513 |
| 32 | Bform | 10.1080/07391102.2002.10506775 |
| 33 | Bform | 10.1002/0471142700.nc0711s11 |
| 34 | Bform | Darwish, Maged A., "Stability of Nucleic Acid Secondary Structures and Their Contribution to Gene Expression" (2010). *Seton Hall University Dissertations and Theses (ETDs)*. 1435. |
| 35 | Bform-DNA | 10.1002/chir.22064 |
| 36 | Bform-DNA | 10.1093/nar/gkl513 |
| 37 | Bform-DNA | 10.1016/j.bpj.2016.11.018 |
| 38 | Bform-DNA | 10.3762/bjoc.14.5 |
| 39 | Bform-DNA | PMID: 6277914 |
| 40 | Bform-DNA | 10.1080/07391102.1989.10506546 |
| 41 | Bform-DNA | 10.1111/j.1751-1097.1986.tb04667.x |
| 42 | Bform-DNA | 10.1002/bip.1979.360180418 |
| 43 | Bform-DNA | 10.1093/nar/15.18.7627 |
| 44 | Bform-DNA | 10.1093/nar/13.13.4983 |
| 45 | Bform-DNA | 10.1002/bip.360231103 |
| 46 | Bform-DNA | 10.1002/bip.1981.360200702 |
| 47 | Bform-DNA | 10.1039/c8ra05179h |
| 48 | Zform-DNA | 10.1073/pnas.94.16.8421 |
| 49 | Zform-DNA | 10.1093/nar/gkp026 |
| 50 | Zform-DNA | 10.1073/pnas.0911528107 |
| 51 | Zform-DNA | 10.1111/j.1747-0285.2009.00847.x |
| 52 | Zform-DNA | 10.1016/0076-6879(95)46006-3 |

| | | |
|---|---|---|
| 53 | Zform-DNA | 10.1371/journal.pone.0198418 |
| 54 | Zform-DNA | 10.1021/jp3085556 |
| 55 | Zform-DNA | 10.1002/anie.201001561 |
| 56 | Zform-DNA | Darwish, Maged A., "Stability of Nucleic Acid Secondary Structures and Their Contribution to Gene Expression" (2010). *Seton Hall University Dissertations and Theses (ETDs)*. 1435 |
| 57 | Zform-DNA | 10.1002/chir.22064 |
| 58 | Zform-DNA | 10.1038/srep09943 |
| 59 | Zform-DNA | 10.3762/bjoc.14.5 |
| 60 | Zform-DNA | PMID: 6277914 |
| 61 | Zform-DNA | 10.1080/07391102.1989.10506546 |
| 62 | Zform-DNA | 10.1111/j.1751-1097.1986.tb04667.x |
| 63 | Zform-DNA | 10.1002/bip.360320307 |
| 64 | Zform-DNA | 10.1093/nar/15.18.7627 |
| 65 | Zform-DNA | 10.1093/nar/13.13.4983 |
| 66 | Zform-DNA | 10.1002/bip.360231103 |
| 67 | Zform-DNA | 10.1093/nar/18.8.2141 |
| 68 | Zform-DNA | 10.1021/bi00316a004 |
| 69 | Antiparallel G quadruplex | 10.3762/bjoc.14.5 |
| 70 | Antiparallel G quadruplex | Darwish, Maged A., "Stability of Nucleic Acid Secondary Structures and Their Contribution to Gene Expression" (2010). *Seton Hall University Dissertations and Theses (ETDs)*. 1435 |
| 71 | Antiparallel G quadruplex | 10.1080/07391102.1989.10506546 |
| 72 | Antiparallel G quadruplex | 10.1002/bip.1979.360180418 |
| 73 | Antiparallel G quadruplex | 10.1002/bip.360290205 |
| 74 | Antiparallel G quadruplex | 10.1002/bip.360271205 |
| 75 | Antiparallel G quadruplex | 10.1002/bip.360231103 |
| 76 | Antiparallel G quadruplex | 10.1038/s41598-017-15797-5 |
| 77 | Antiparallel G quadruplex | 10.1186/1471-2164-15-1032 |

| | | |
|---|---|---|
| 78 | Antiparallel G quadruplex | 10.1007/s00249-018-1312-4 |
| 79 | Antiparallel G quadruplex | *10.1093/nar/gkw970* |
| 80 | Antiparallel G quadruplex | 10.1039/b915955j |
| 81 | Antiparallel G quadruplex | 10.1016/j.tig.2018.11.001c |
| 82 | Antiparallel G quadruplex | *10.1093/nar/gky902* |
| 83 | Antiparallel G quadruplex | 10.1002/cpnc.23 |
| 84 | Antiparallel G quadruplex | 10.1021/acs.jchemed.7b00160 |
| 85 | Antiparallel G quadruplex | 10.5114/bta.2012.46592 |
| 86 | Antiparallel G quadruplex | 10.3390/molecules24101863 |
| 87 | Antiparallel G quadruplex | 10.1155/2017/9170371 |
| 88 | Antiparallel G quadruplex | 10.1074/jbc.M117.776211 |
| 89 | Antiparallel G quadruplex | 10.1093/nar/gkl348 |
| 90 | Antiparallel G quadruplex | 10.1186/1471-2164-10-362 |
| 91 | Hybrid G quadruplex | 10.1002/anie.201709184 |
| 92 | Hybrid G quadruplex | *10.1093/nar/gky757* |
| 93 | Hybrid G quadruplex | 10.1093/nar/gky757 |
| 94 | Hybrid G quadruplex | 10.1038/s41598-018-20852-w |
| 95 | Hybrid G quadruplex | 10.3389/fchem.2018.00281 |
| 96 | Hybrid G quadruplex | *10.1093/nar/gkw970* |
| 97 | Hybrid G quadruplex | 10.1002/cpnc.23 |
| 98 | Hybrid G quadruplex | 10.1021/acs.jchemed.7b00160 |
| 99 | Hybrid G quadruplex | 10.1039/c8cp04728f |
| 100 | Hybrid G quadruplex | 10.1155/2017/9170371 |
| 101 | Hybrid G quadruplex | 10.1074/jbc.M117.776211 |
| 102 | Hybrid G quadruplex | 10.1093/nar/gkl348 |
| 103 | Hybrid G quadruplex | 10.1186/1471-2164-10-362 |
| 104 | Parallel G quadruplex | 10.1016/j.ymeth.2007.02.009 |
| 105 | Parallel G quadruplex | 10.1093/nar/gkp026 |
| 106 | Parallel G quadruplex | 10.1002/anie.201709184 |
| 107 | Parallel G quadruplex | *10.1002/bip.10112* |
| 108 | Parallel G quadruplex | 10.5114/bta.2012.46592 |

| | | |
|---|---|---|
| 109 | Parallel G quadruplex | 10.1002/rcm.778 |
| 110 | Parallel G quadruplex | 10.1002/0471142700.nc0711s11 |
| 111 | Parallel G quadruplex | *10.1093/nar/gky757* |
| 112 | Parallel G quadruplex | 10.1093/nar/gky757 |
| 113 | Parallel G quadruplex | 10.1038/s41598-018-20852-w |
| 114 | P-g4 | 10.1038/srep09255 |
| 115 | Parallel G quadruplex | 10.1186/1471-2164-15-1032 |
| 116 | Parallel G quadruplex | *10.1093/nar/gkw970* |
| 117 | Parallel G quadruplex | *10.1093/nar/gkx100* |
| 118 | Parallel G quadruplex | 10.1039/b915955j |
| 119 | Parallel G quadruplex | *10.1093/nar/gkv266* |
| 120 | Parallel G quadruplex | 10.1016/j.tig.2018.11.001c |
| 121 | Parallel G quadruplex | *10.1093/nar/gky902* |
| 122 | Parallel G quadruplex | 10.1002/cpnc.23. |
| 123 | Parallel G quadruplex | 10.1021/acs.jchemed.7b00160 |
| 124 | Parallel G quadruplex | 10.5114/bta.2012.46592 |
| 125 | Parallel G quadruplex | 10.1039/c8cp04728f |
| 126 | Parallel G quadruplex | 10.1042/BSR20171128 |
| 127 | Parallel G quadruplex | 10.1101/gad.305862.117 |
| 128 | Parallel G quadruplex | 10.1074/jbc.M117.776211 |
| 129 | Parallel G quadruplex | 10.7554/eLife.26884 |
| 130 | Antiparallel triplex | 10.1093/nar/gks410 |
| 131 | Antiparallel triplex | 10.1093/nar/20.15.3859 |
| 132 | Antiparallel triplex | 10.1093/nar/14.24.10071 |
| 133 | Antiparallel triplex | 10.1016/s0301-4622(00)00200-3 |
| 134 | Antiparallel triplex | 10.1093/nar/gkv496 |
| 135 | Antiparallel triplex | 10.1093/nar/28.5.1162 |
| 136 | Antiparallel triplex | 10.1021/bi301381 |
| 137 | Antiparallel triplex | 10.1021/acs.jpcb.7b07591 |
| 138 | Antiparallel triplex | *10.1093/nar/gkv1017* |

| | | |
|---|---|---|
| 139 | Antiparallel triplex | 10.1021/ic5017565 |
| 140 | Antiparallel triplex | 10.1080/07391102.2015.1077344 |
| 141 | Antiparallel triplex | 10.1016/j.ijbiomac.2017.05.115 |
| 142 | Antiparallel triplex | 10.3390/ijms17020211 |
| 143 | Antiparallel triplex | *10.1093/nar/gkx100* |
| 144 | Antiparallel triplex | 10.1039/c5ob00535c |
| 145 | Antiparallel triplex | 10.1038/s41598-017-15797-5 |
| 146 | Antiparallel triplex | 10.1016/j.abb.2017.11.008 |
| 147 | Antiparallel triplex | 10.1038/s41598-018-31387-5 |
| 148 | Antiparallel triplex | 10.1371/journal.pone.0037939 |
| 149 | Antiparallel triplex | 10.1080/07391102.2016.1160257 |
| 150 | Parallel triplex | 10.1016/0076-6879(95)46005-5 |
| 151 | Parallel triplex | 10.1021/bi025937m |
| 152 | Parallel triplex | 10.1021/bi0004201 |
| 153 | Parallel triplex | 10.1093/nar/27.16.3371 |
| 154 | Parallel triplex | 10.1042/BJ20141023 |
| 155 | Parallel triplex | 10.1016/s0301-4622(02)00380-0 |
| 156 | Parallel triplex | 10.1021/bi010666l |
| 157 | Parallel triplex | 10.1002/jms.1277 |
| 158 | Parallel triplex | 10.1002/jccs.201000072 |
| 159 | Parallel triplex | 10.1016/s0162-0134(02)00444-0 |
| 160 | Parallel triplex | 10.1093/nar/18.12.3557 |
| 161 | Parallel triplex | 10.1093/nar/28.5.1162 |
| 162 | Parallel triplex | 10.1021/bi301381 |
| 163 | Parallel triplex | 10.1021/acs.jpcb.7b07591 |
| 164 | Parallel triplex | *10.1093/nar/gkv1017* |
| 165 | Parallel triplex | 10.1021/ic5017565 |
| 166 | Parallel triplex | 10.1080/07391102.2015.1077344 |
| 167 | Parallel triplex | 10.1016/j.ijbiomac.2017.05.115 |
| 168 | Parallel triplex | 10.3390/ijms17020211 |

| | | |
|---|---|---|
| 169 | Parallel triplex | *10.1093/nar/gkx100* |
| 170 | i motif | 10.1093/nar/gkn380 |
| 171 | i motif | *10.1093/nar/gky390* |
| 172 | i motif | *10.1093/nar/gkw402* |
| 173 | i motif | 10.1166/jnn.2010.2848 |
| 174 | i motif | 10.3389/fchem.2018.00281 |
| 175 | i motif | 10.1039/c8ra05179h |
| 176 | i motif | 10.1039/b919600e |
| 177 | i motif | 10.1002/cbic.201500182 |
| 178 | i motif | 10.1166/jnn.2010.2848 |
| 179 | i motif | *10.1093/nar/gky390* |
| 180 | i motif | 10.1039/c5cc05111h |
| 181 | i motif | 10.1039/c6cp04418b |
| 182 | i motif | *10.1093/nar/gky390* |
| 183 | i motif | 10.1039/c7cp06235d |
| 184 | i motif | 10.1039/c3cc46594b |
| 185 | i motif | 10.1039/c7cp05380k |
| 186 | i motif | *10.1093/nar/gky035* |
| 187 | i motif | 10.1038/s41557-018-0046-3 |
| 188 | i motif | 10.1093/nar/gkn380 |
| 189 | i motif | *10.1093/nar/gkx090* |
| 190 | i motif | 10.1039/c4cc06824f |
| 191 | i motif | *10.1093/nar/gkz046* |
| 192 | i motif | *10.1093/nar/gkw402* |
| 193 | i motif | 10.1021/acsomega.9b00784 |
| 194 | i motif | 10.1021/acs.biochem.7b00628 |
| 195 | i motif | 10.3390/nano8040226 |
| 196 | i motif | 10.1039/c8ob00920a |
| 197 | Parallel duplex | 10.1016/j.ymeth.2012.03.011 |
| 198 | Parallel duplex | 10.1093/nar/16.14.6659 |
| 199 | Parallel duplex | 10.1021/acs.langmuir.6b03253 |

| | | |
|---|---|---|
| 200 | Parallel duplex | 10.1093/nar/gkp133 |
| 201 | Parallel duplex | 10.1002/chir.22064 |
| 202 | Parallel duplex | 10.1021/bi010666 |
| 203 | DNA-RNA duplex | 10.1002/chir.22064 |
| 204 | DNA-RNA duplex | 10.1371/journal.pone.0143354 |
| 205 | DNA-RNA duplex | 10.1039/c0mb00258e |
| 206 | DNA stemloop | Darwish, Maged A., "Stability of Nucleic Acid Secondary Structures and Their Contribution to Gene Expression" (2010). *Seton Hall University Dissertations and Theses (ETDs)*. 1435 |
| 207 | RNA stemloop | Darwish, Maged A., "Stability of Nucleic Acid Secondary Structures and Their Contribution to Gene Expression" (2010). *Seton Hall University Dissertations and Theses (ETDs)*. 1435 |
| 208 | Antiparallel triplex | 10.3390/ijms17020211 |
| 209 | G triplex | 10.1039/c8an01208c |
| 210 | G triplex | 10.1042/BJ20141023 |
| 211 | G triplex | *10.1093/nar/gku1084* |
| 212 | iG Pentaplex | *10.1093/nar/gkv266* |
| 213 | iG Quadruplex | *10.1093/nar/gkv266* |

**Table S2. Superimposed training CD spectrum corresponding to 16 different nucleic acid secondary structures. Note that (e-g), (i-j), (l-n) and (p-q) correspond to different forms of hybrid G-quadruplex, i-motif, parallel triplex and anti-parallel triplex.**

(a) Z form

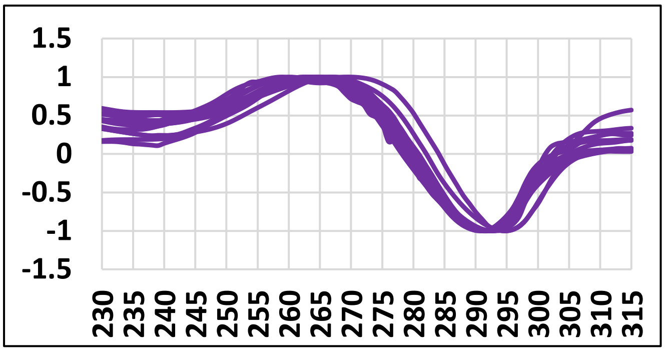


(b) B form

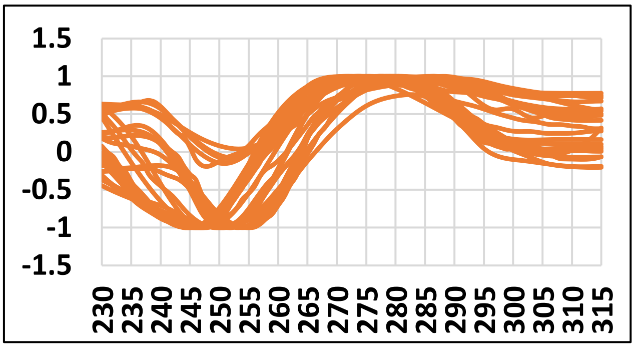


(c) Parallel G quadruplex

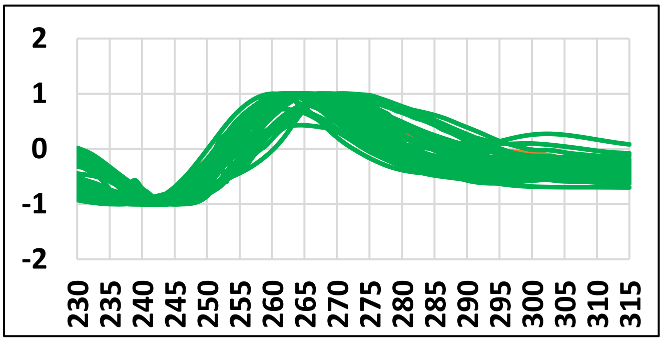


(d) Antiparallel G quadruplex

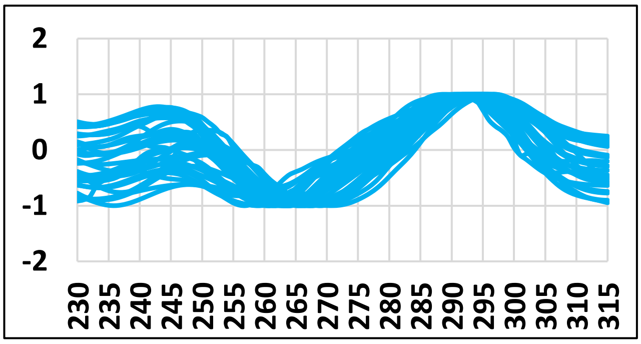


(e) Hybrid G quadruplex

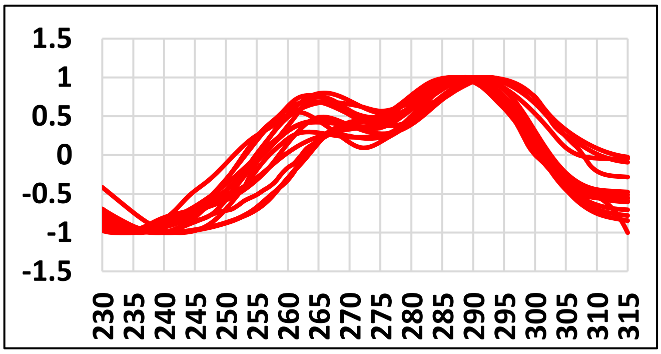


(f) Hybrid G quadruplex

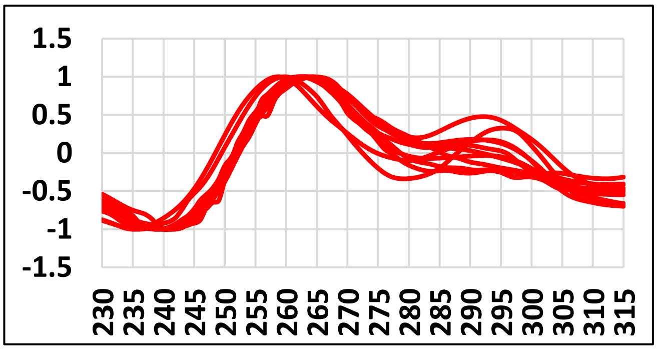

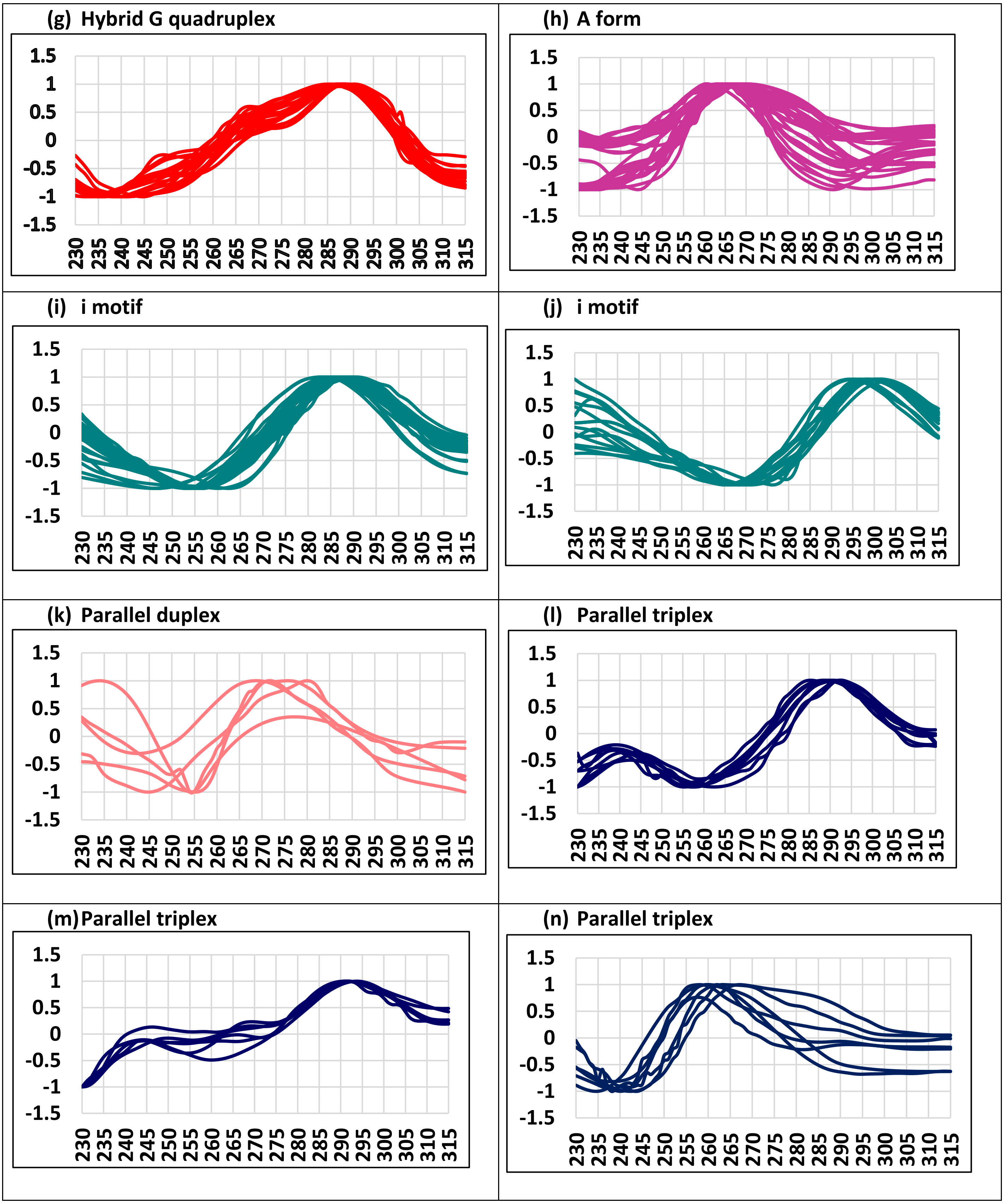
(g) Hybrid G quadruplex
(h) A form
(i) i motif
(j) i motif
(k) Parallel duplex
(l) Parallel triplex
(m) Parallel triplex
(n) Parallel triplex
1.5
1
0.5
0
-0.5
-1
-1.5
230 235 240 245 250 255 260 265 270 275 280 285 290 295 300 305 310 315

(o) RNA stemloop

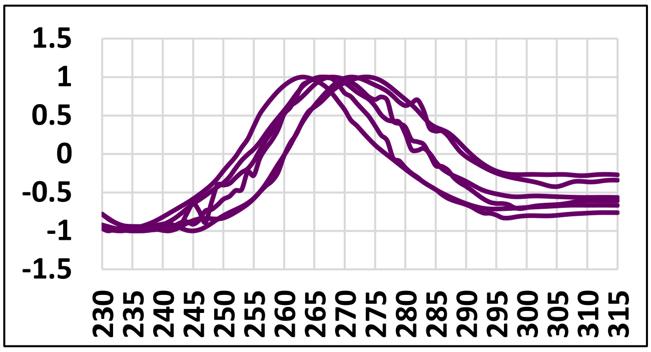


(p) Antiparallel triplex

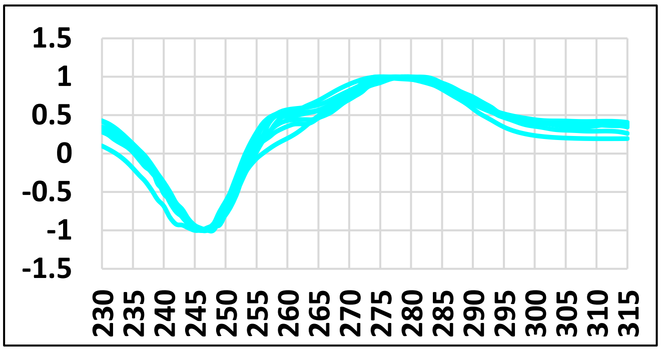


(q) Antiparallel triplex

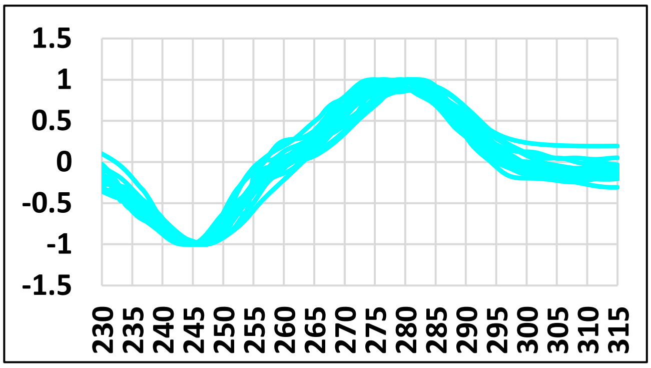


(r) DNA RNA duplex

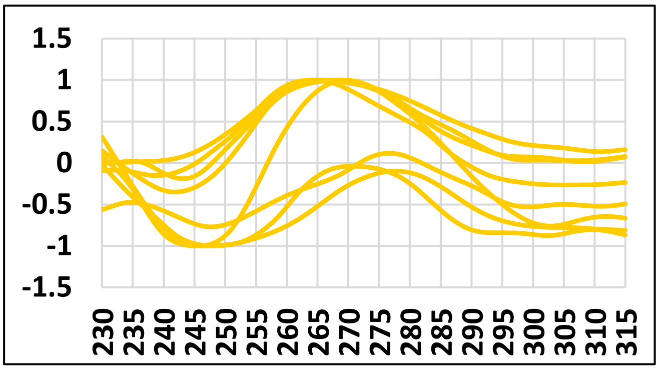


(s) DNA stemloop

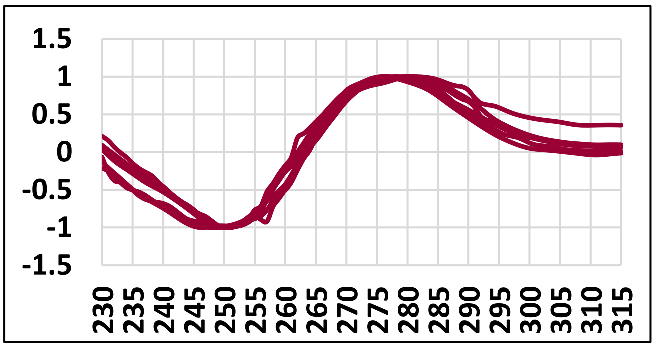


(x) G triplex

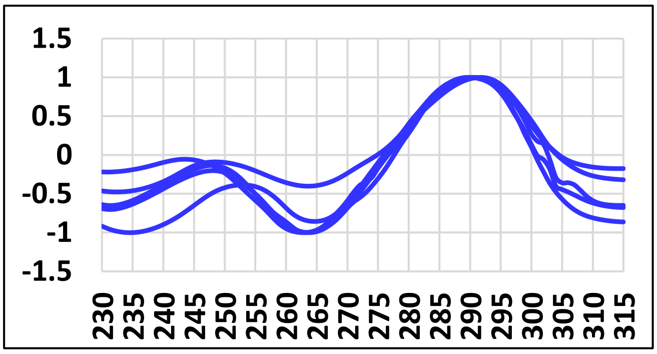


(y) iG pentaplex

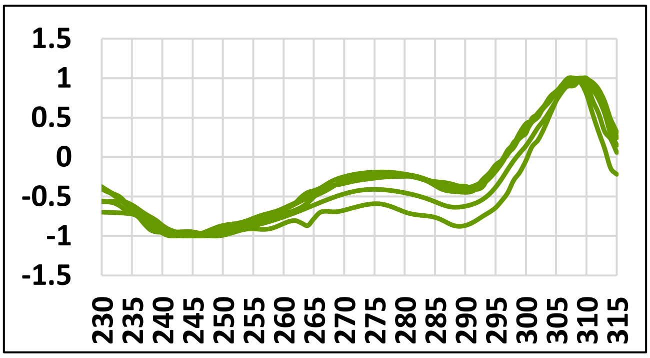


(z) iG quadruplex

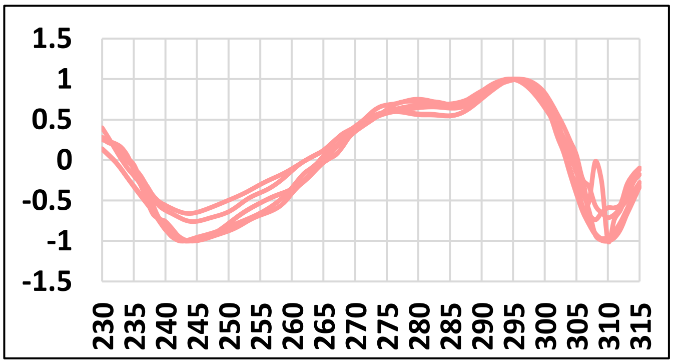

**Table S3**. **The 16*16 confusion matrix corresponding to *XGBoost* (brown colored cell), *nnet* (grey colored cell) and *Kohonen* (green colored cell) algorithms.** The total number of accurate and false predictions for the 242 test datasets (**Figure 1c**) is indicated under 16 different nucleic acids conformations. Note that the 16x16 matrix corresponds to the 16 known (1st column) and the predicted (1st row) secondary structures. (Note: APT-Antiparallel triplex, A G4 – Antiparallel G quadruplex, H G4 - Hybrid G quadruplex, IM – i-motif, PD – parallel duplex, P G4 – Parallel G quadruplex, PT – Parallel triplex)

| Nucleic acid form | | Aform | AP G4 | A PT | Bform | DNA stemloop | DNA-RNA duplex | G triplex | H G4 | iG Pentaplex | iG Quadruplex | I M | P D | P G4 | P T | RNA stemloop | Zform |
|---|---|---|---|---|---|---|---|---|---|---|---|---|---|---|---|---|---|
| **Aform** | *XGBoost* | 23 | 0 | 0 | 0 | 0 | 0 | 0 | 0 | 0 | 0 | 0 | 0 | 0 | 0 | 0 | 0 |
| | *nnet* | 11 | 0 | 0 | 0 | 9 | 0 | 0 | 0 | 0 | 0 | 0 | 0 | 1 | 1 | 0 | 1 |
| | *Kohonen* | 13 | 0 | 2 | 0 | 0 | 6 | 1 | 0 | 0 | 0 | 0 | 1 | 0 | 0 | 0 | 0 |
| **AP G4** | *XGBoost* | 0 | 26 | 0 | 0 | 0 | 0 | 0 | 0 | 0 | 0 | 0 | 0 | 0 | 0 | 0 | 0 |
| | *nnet* | 0 | 23 | 0 | 0 | 0 | 0 | 0 | 1 | 0 | 0 | 1 | 0 | 0 | 1 | 0 | 0 |
| | *Kohonen* | 0 | 17 | 0 | 0 | 0 | 0 | 0 | 4 | 2 | 0 | 3 | 0 | 0 | 0 | 0 | 0 |
| **APT** | *XGBoost* | 0 | 0 | 28 | 0 | 0 | 0 | 0 | 0 | 0 | 0 | 0 | 0 | 0 | 0 | 0 | 0 |
| | *nnet* | 0 | 0 | 26 | 0 | 0 | 0 | 0 | 0 | 0 | 0 | 2 | 0 | 0 | 0 | 0 | 0 |
| | *Kohonen* | 0 | 0 | 12 | 2 | 1 | 2 | 2 | 0 | 0 | 0 | 0 | 3 | 1 | 0 | 0 | 0 |
| **Bform** | *XGBoost* | 0 | 0 | 0 | 18 | 0 | 0 | 0 | 0 | 0 | 0 | 0 | 0 | 1 | 0 | 0 | 0 |
| | *nnet* | 0 | 0 | 4 | 7 | 1 | 1 | 0 | 1 | 0 | 0 | 1 | 0 | 1 | 0 | 0 | 2 |
| | *Kohonen* | 1 | 0 | 2 | 10 | 2 | 1 | 1 | 0 | 0 | 0 | 0 | 0 | 2 | 0 | 0 | 0 |
| **DNA stemloop** | *XGBoost* | 0 | 0 | 0 | 0 | 3 | 0 | 0 | 0 | 0 | 0 | 0 | 0 | 0 | 0 | 0 | 0 |
| | *nnet* | 0 | 0 | 0 | 0 | 3 | 0 | 0 | 0 | 0 | 0 | 0 | 0 | 0 | 0 | 0 | 0 |
| | *Kohonen* | 0 | 0 | 0 | 0 | 2 | 1 | 0 | 0 | 0 | 0 | 0 | 0 | 0 | 0 | 0 | 0 |
| **DNA RNA duplex** | *XGBoost* | 0 | 0 | 0 | 0 | 0 | 5 | 0 | 0 | 0 | 0 | 0 | 0 | 0 | 0 | 0 | 0 |
| | *nnet* | 1 | 0 | 0 | 1 | 0 | 2 | 0 | 0 | 0 | 0 | 0 | 0 | 1 | 0 | 0 | 1 |
| | *Kohonen* | 2 | 0 | 0 | 0 | 0 | 0 | 0 | 0 | 0 | 0 | 1 | 2 | 0 | 0 | 0 | 0 |
| **G triplex** | *XGBoost* | 0 | 0 | 0 | 0 | 0 | 0 | 5 | 0 | 0 | 0 | 0 | 0 | 0 | 0 | 0 | 0 |
| | *nnet* | 0 | 0 | 0 | 0 | 0 | 0 | 4 | 0 | 0 | 0 | 0 | 0 | 1 | 0 | 0 | 0 |
| | *Kohonen* | 0 | 0 | 0 | 0 | 0 | 0 | 3 | 1 | 0 | 0 | 0 | 0 | 1 | 0 | 0 | 0 |

| | | | | | | | | | | | | | | | | | |
|---|---|---|---|---|---|---|---|---|---|---|---|---|---|---|---|---|---|
| **H G4** | *XGBoost* | 0 | 0 | 0 | 0 | 0 | 0 | 0 | 26 | 0 | 0 | 0 | 0 | 0 | 0 | 0 | 0 |
| | *nnet* | 0 | 1 | 0 | 0 | 0 | 0 | 0 | 25 | 0 | 0 | 0 | 0 | 0 | 0 | 0 | 0 |
| | *Kohonen* | 1 | 1 | 0 | 0 | 0 | 0 | 0 | 16 | 0 | 0 | 3 | 0 | 5 | 0 | 0 | 0 |
| **Ig pentaplex** | *XGBoost* | 0 | 0 | 0 | 0 | 0 | 0 | 0 | 0 | 3 | 0 | 0 | 0 | 0 | 0 | 0 | 0 |
| | *nnet* | 0 | 0 | 0 | 0 | 0 | 0 | 0 | 0 | 3 | 0 | 0 | 0 | 0 | 0 | 0 | 0 |
| | *Kohonen* | 0 | 0 | 0 | 0 | 0 | 0 | 0 | 0 | 3 | 0 | 0 | 0 | 0 | 0 | 0 | 0 |
| **Ig quadruplex** | *XGBoost* | 0 | 0 | 0 | 0 | 0 | 0 | 0 | 0 | 0 | 3 | 0 | 0 | 0 | 0 | 0 | 0 |
| | *nnet* | 0 | 0 | 0 | 0 | 0 | 0 | 0 | 0 | 0 | 3 | 0 | 0 | 0 | 0 | 0 | 0 |
| | *Kohonen* | 0 | 0 | 0 | 0 | 0 | 0 | 0 | 0 | 0 | 3 | 0 | 0 | 0 | 0 | 0 | 0 |
| **IM** | *XGBoost* | 0 | 0 | 0 | 0 | 0 | 0 | 0 | 0 | 0 | 0 | 40 | 0 | 0 | 0 | 0 | 0 |
| | *nnet* | 0 | 0 | 0 | 0 | 0 | 0 | 0 | 0 | 0 | 0 | 40 | 0 | 0 | 0 | 0 | 0 |
| | *Kohonen* | 0 | 0 | 0 | 0 | 0 | 1 | 0 | 0 | 0 | 0 | 38 | 1 | 0 | 0 | 0 | 0 |
| **PD** | *XGBoost* | 0 | 0 | 0 | 0 | 0 | 0 | 0 | 0 | 0 | 0 | 0 | 5 | 0 | 0 | 0 | 0 |
| | *nnet* | 0 | 0 | 0 | 1 | 0 | 0 | 0 | 0 | 0 | 0 | 0 | 4 | 0 | 0 | 0 | 0 |
| | *Kohonen* | 0 | 0 | 3 | 0 | 0 | 1 | 1 | 0 | 0 | 0 | 0 | 0 | 0 | 0 | 0 | 0 |
| **P G4** | *XGBoost* | 0 | 0 | 0 | 0 | 0 | 0 | 0 | 0 | 0 | 0 | 1 | 0 | 18 | 0 | 0 | 0 |
| | *nnet* | 0 | 0 | 0 | 0 | 0 | 1 | 0 | 0 | 0 | 0 | 1 | 0 | 14 | 0 | 0 | 1 |
| | *Kohonen* | 0 | 0 | 2 | 0 | 0 | 0 | 1 | 0 | 0 | 0 | 1 | 0 | 15 | 0 | 0 | 0 |
| **PT** | *XGBoost* | 0 | 0 | 0 | 0 | 0 | 0 | 0 | 0 | 0 | 0 | 0 | 0 | 0 | 16 | 0 | 0 |
| | *nnet* | 0 | 0 | 1 | 0 | 0 | 0 | 0 | 0 | 0 | 0 | 0 | 0 | 0 | 15 | 0 | 0 |
| | *Kohonen* | 2 | 0 | 3 | 0 | 0 | 0 | 0 | 0 | 0 | 0 | 0 | 1 | 3 | 7 | 0 | 0 |
| **RNA stemloop** | *XGBoost* | 0 | 0 | 0 | 0 | 0 | 0 | 0 | 0 | 0 | 0 | 0 | 0 | 0 | 0 | 3 | 0 |
| | *nnet* | 0 | 0 | 0 | 0 | 0 | 0 | 0 | 0 | 0 | 0 | 0 | 0 | 0 | 0 | 3 | 0 |
| | *Kohonen* | 0 | 0 | 0 | 0 | 0 | 0 | 0 | 0 | 0 | 0 | 0 | 1 | 0 | 0 | 2 | 0 |
| **Zform** | *XGBoost* | 0 | 0 | 0 | 0 | 0 | 0 | 0 | 0 | 0 | 0 | 0 | 0 | 0 | 0 | 0 | 18 |
| | *nnet* | 1 | 0 | 0 | 0 | 0 | 0 | 0 | 0 | 0 | 0 | 1 | 0 | 0 | 0 | 0 | 16 |
| | *Kohonen* | 2 | 0 | 0 | 1 | 0 | 0 | 0 | 0 | 0 | 0 | 1 | 0 | 0 | 0 | 0 | 14 |

**Table S4.** Summary of the secondary structural prediction from CD spectra using *XGBoost*, *nnet* and *Kohonen* algorithms. Note that the prediction accuracy is higher for *XGBoost* algorithm compared to the others.

| Nucleic acid form | Total test set | *XGBoost* | *nnet* | *Kohonen* |
|---|---|---|---|---|
| Aform | 23 | 23 | 11 | 13 |
| Antiparallel G-Quadruplex | 26 | 26 | 23 | 17 |
| Antiparallel triplex | 28 | 28 | 26 | 12 |
| Bform | 19 | 18 | 7 | 10 |
| DNA stemloop | 3 | 3 | 3 | 2 |
| DNA-RNA duplex | 5 | 5 | 2 | 0 |
| G triplex | 5 | 5 | 4 | 3 |
| Hybrid G-Quadruplex | 26 | 26 | 25 | 16 |
| iG Pentaplex | 3 | 3 | 3 | 3 |
| iG Quadruplex | 3 | 3 | 3 | 3 |
| i-motif | 40 | 40 | 40 | 38 |
| Parallel duplex | 5 | 5 | 4 | 0 |
| Parallel G-Quadruplex | 19 | 18 | 14 | 15 |
| Parallel triplex | 16 | 16 | 15 | 7 |
| RNA stemloop | 3 | 3 | 3 | 2 |
| Zform | 18 | 18 | 16 | 14 |
| Total | 242 | 240 | 199 | 155 |
| Accuracy | | 99% | 82% | 64% |
